# Determinants of fleshiness or dryness for fruits at maturity: fruit shapes and pericarp permeability

**DOI:** 10.1101/2025.05.13.653672

**Authors:** Shunli Yu, Canran Liu, Hui Liu, Xiuli Gao, Carol C. Baskin

**Affiliations:** Key Laboratory of Vegetation and Environmental Changes, Institute of Botany, Chinese Academy of Sciences, Beijing, 100093, China; Arthur Rylah Institute for Environmental Research, Department of Energy, Environment and Climate Action, Heidelberg, VIC 3084, Australia; South China Botanical Garden, Chinese Academy of Sciences, Guangzhou, 510650, China; University of Chinese Academy of Sciences, Beijing, 100049, China; Department of Biology, University of Kentucky, Lexington, Kentucky, USA

**Keywords:** fleshy and dry fruits, fruit type diversity, fruit water content variation, modes of water loss in fruits, open and shaded habitats, propagule shape diversity, waxy fruit epidermis

## Abstract

[A hypothesis was proposed that fleshiness or dryness of fruits at maturity were determined just by fruit shapes and pericarp permeability influenced by heterogeneous habitats] The ecological origins of diverse fruit types still remain unresolved due to the lack of knowledge about fruit water content variation patterns during fruit development and after fruit maturity. In this study, we explore factors influencing fleshiness or dryness of fruits at maturity through examining the water content variation patterns before and after fruit maturity, and identify biological and environmental mechanisms about fleshiness or not for fruits at maturity, using data from a survey on fruit shapes of 29760 species and one experiment involving water content measure of fruits. Results show that water content in majority of the fleshy (except for drupes and rosehips) and dry fruits increases or decreases, respectively, during fruit development i.e. prior to maturity, and the water variation patterns are determined by fruit shapes and pericarp permeability. After maturation, water loss occurs slowly at room temperature for fleshy fruits, but it is fast for most dry fruits. In general, declining fruit water content due to pericarp lignification occurring and rising pericarp permeability with the development of dry fruits in later stage of fruit development as well as water imbalance between supply and loss induced by open habitats may lead to the formation of dryness for elongated or flaky dry fruits owing to allometric growth between pedicel diameter (or fruit mass) and fruit surface area. Resulted from water balance, the emergence of fleshiness in mature fleshy fruits is attributed to slow water loss through the pericarp influenced by globular shapes (i.e. greater fruit thickness) and pericarp waxification induced by shaded habitats, or by sporoderm lignification occurring (inside the pericarp) induced by open habitats for a subset of fleshy fruits such as drupes and rosehips.

## Introduction

Numerous studies have explored the biological characteristics of fleshy and dry fruits (1,2) and their ecological relationships with environmental factors (3–5). However, much remains to be learned about the essential difference and mechanisms in fruit water content dynamics between fleshy and dry fruits, especially in their water adaptation strategies to heterogeneous habitats. It is known that dry fruits, with a low water content when ripe, typically have a period of higher water content during fruit development (6) and fleshy fruits are defined as a kind of fruits in which the pericarp and accessory parts develop into succulent tissues at maturity. However, we still know little about water content dynamics of the two fruit types during their development and after mature. Inexact knowledge of water physiology in fleshy and dry fruits hinders the solution of determinants of fleshiness or dryness for fruits at maturity in a long term.

Selection pressures that shape fruit traits, especially in fruit fleshiness or dryness, have puzzled ecologist and evolutionary biologist for several decades (7–9), yet they remain subjects of considerable controversy due to lack of knowledge in fruit water dynamics, shapes and pericarp traits (10). Previous studies have stated that plant-animal interactions, correlated evolution of internal organs of plants and environmental adaptations might lay a strong foundation for understanding the plant propagule traits in fleshiness or dryness (10–11). However, the factors determining whether fruit traits are fleshy or dry, and the extent to which they are shaped by biotic factors such as dispersers or internal structure of plants, remains unclear (12). Exploring the essential variation rules in propagule feature in relation to water ecophysiology for fruit types like fleshy and dry fruits can provide a knowledge basis for addressing co-evolution of internal organs of plants and environmental adaptation. Therefore, we conducted experiments to compare the water dynamics of various fruit types before and after fruit ripening and to identify whether there are significant variations in water content dynamics between fleshy and dry fruits. Additionally, it is speculated that the formation of fleshiness in fleshy fruits may result from water balance, however, exact cause remains a mystery. Transpiration, photosynthesis and the integration of structural components of fruits are the main final destination of water transported from the pedicels and twigs. Barrier properties of plant cuticles in pericarp have been proved to play a vital role in protecting against water loss (13). However, whether there are allometric growth relationships or trade-off in water supply and loss between fruits and pedicel or whether fleshiness of fruits is related to other plant traits such as fruit shapes and pericarp traits needs to be investigated.

Fleshy and dry fruits, two inclusive characteristics with huge phylogenetic differentiation, evolved through concerted convergence, undergoing respectively concurrent adaptations to shaded and open habitats (7–8, 14). However, the strongest evidence and evolution momentum for concerted convergence in fruit types have yet to be uncovered (8). The distribution of fleshy-fruited species and their close correlation with shaded habitats suggest that they may originate in the closed environment or forest understories (Fig. 1) (15, 16), where solar radiation, air temperature and air flow are relatively lower or weaker compared to the open habitats. Therefore, the origin causation of fleshy/dry fruits is more likely associated with higher/lower water availability related to environmental factors (17). Soil moisture was not found to be significantly related to the distribution of fleshy-fruited species (15, 18), but temperature and rainfall amount were (5). High water availability and high photosynthate allocation may lead to the emergence of fleshy fruits with larger seeds (17). However, we know little about the relative importance of abiotic factors, such as light, temperature and wind, in determining diverse fruit types. The details of how fleshiness or dryness came into being for ripening fruits and the specifics of their emergence process remain a mystery.

**Fig. 1.**
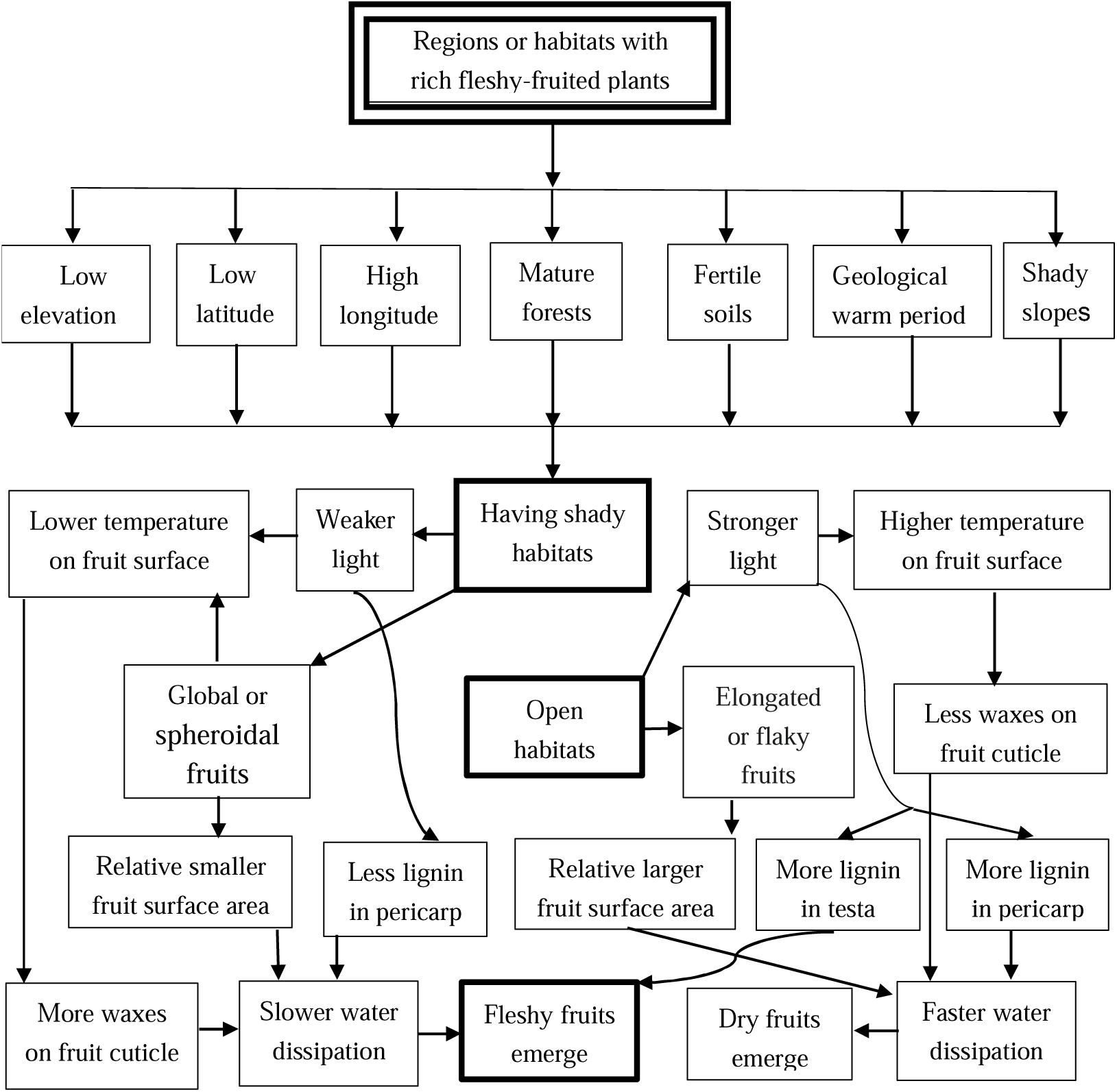
Formation process of fleshy and dry fruits

With regard to fruit shape, a sphere is a very economical shape for packaging seeds (19) and favors water retention due to its minimal surface area for a given weight compared to other shapes. Therefore, extensive studies on the fruit shapes of wild plants can help us understand the adaptation strategies of fruits in relation to their environment. Some of the earliest fossil angiosperm fruits have been found to be approximately globular (20). It is likely that fruit morphology is a result of long-term adaptation and evolution in response to heterogenous environments rather than randomly formed (21), of course, that’s another topic that worth further studying in the future.

In this study, we explore biological and environmental mechanisms about development of fleshy and dry fruits, we determine the differences between fleshy and dry fruits in terms of fruit water content dynamics during fruit development and after fruit maturation. Also, we test which biotic or abiotic factors influence the patterns of these differences in fruit traits. In addition, owing to the relationship between fruit shapes and the rate of water loss, we examine fruit shape variation and explore its correlation with fruit types for wild seed plants distributed in China. The purpose of this study was to explore ecological determinants of fleshiness and dryness for fleshy and dry fruits at maturity through studying water dynamics of different fruit types, combined with fruit shape analysis, pericarp permeability measurement and previous results about the relationships between fleshy-fruited species distribution and shaded habitats ((summarized in 15). In this study, we proposed a hypothesis that diverse water content variation patterns, various fruit shapes and pericarp permeability, influenced by shaded and open habitats (15), may lead to fruit type diversity. We aimed to uncover the biological and environmental mechanism about fleshiness or dryness for fruits at maturity through insight on allometric growth relationships among internal organs of plants and their water adaptation strategies to different habitats.

## Results

### 3.1 Fruit types and shapes

For the two traits (fruit types and fruit shapes), the phylogenetic signal delta values (δ) calculated with the resampled observed data are much larger than those calculated with the randomized data (Fig. 2a-b, Fig. s1). The D values are clearly smaller than 0 for both sample sizes (Fig. 2c). These results clearly demonstrate the presence of a phylogenetic signal in the two traits. Globular or spheroidal fruits show the highest probability being fleshy, followed by small globular fruits, while elongated and flaky fruits show the lowest probability (Fig. 2d).

**Fig. 2.**
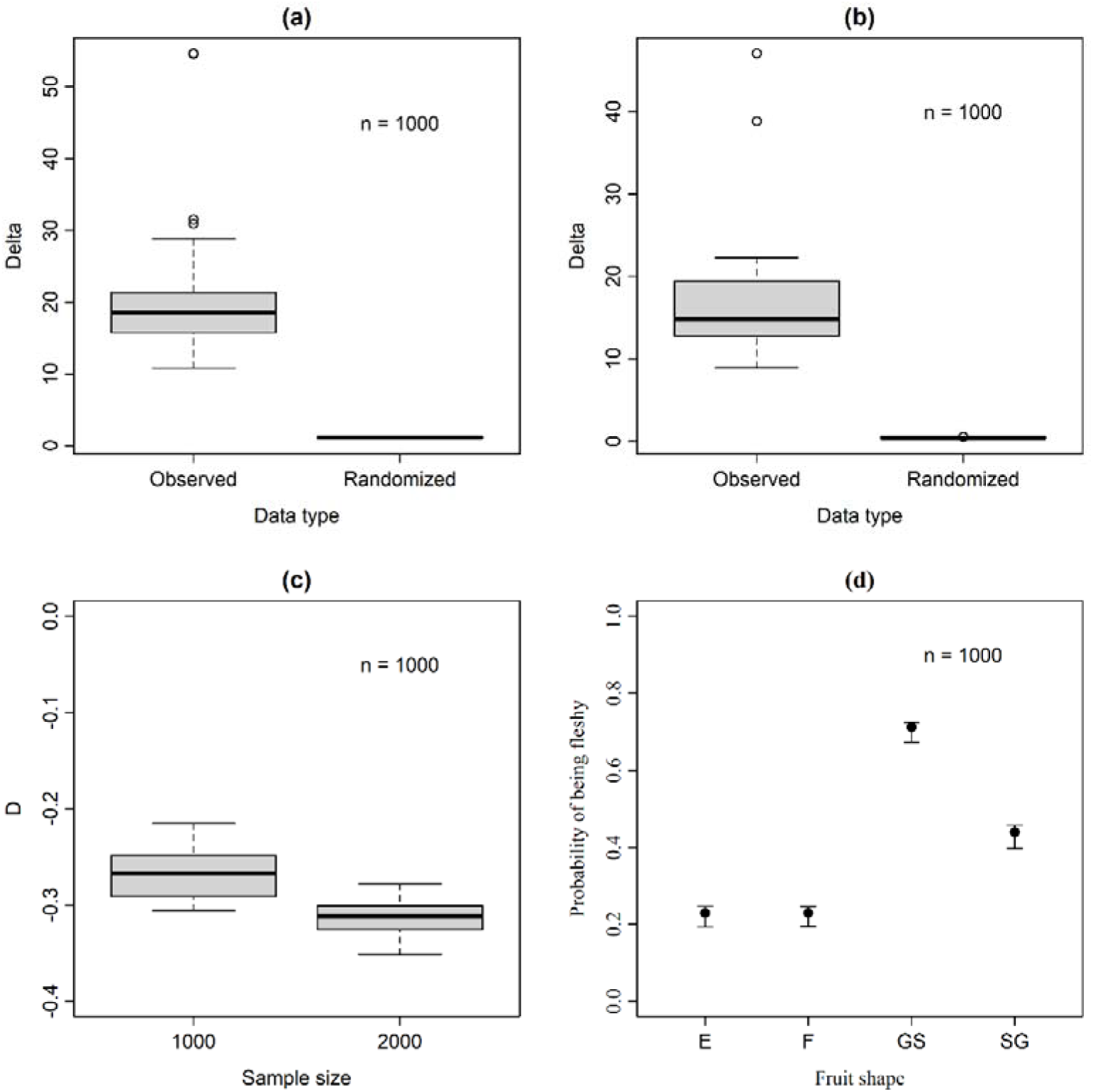
(a) Delta index measuring phylogenetic signal in fruit types. Thirty sets of 1000 species were randomly selected from all the 29760 species. For each set, a value of delta (denoted as observed here) was calculated, and the trait values were randomized and another value of delta (denoted as Random here) was calculated. (b) Delta index measuring phylogenetic signal in fruit shapes. Thirty sets of 1000 species were randomly selected from all the 29760 species. For each set, a value of delta (denoted as Observed here) was calculated, and the trait values were randomized and another value of delta (denoted as Random here) was calculated. (c) D index measuring phylogenetic signal in fruit types. Thirty sets of 1000 and 2000 species were randomly selected from all the 29760 species. For each set, a value of D was calculated. (d) The probability (mean value and 95% confidence interval) that a species has a fleshy fruit for different fruit shapes. Four categories are globular or spheroidal (GS), elongated (E), flaky (F), and small globular (SG).

### 3.2 Water content shift patterns of fruits from juvenile to maturity

Based on fruit water content data throughout the fruit development process, water content of dry fruits decreases generally, while that of most fleshy fruits (except for drupes and rosehips) increases slightly with fruit maturation (Fig. 3). Plant species (belonging to sciophytes) with berries, pomes, pepos, hesperidium and aggregate fruits belong to this type, for example, fruit water content of *Lonicera fragrantissima* is 87.60±1.92% before maturation and 91.04±4.30% at maturation. Water content of drupes and rosehips decreases slightly with fruit maturation, for instance, fruit water content of *Prunus trilobata* is 81.30±1.20% before maturation and 58.95±4.83% at maturation. On the whole, fleshy fruits had a higher water content even at fruit maturity than dry fruits.

**Fig. 3.**
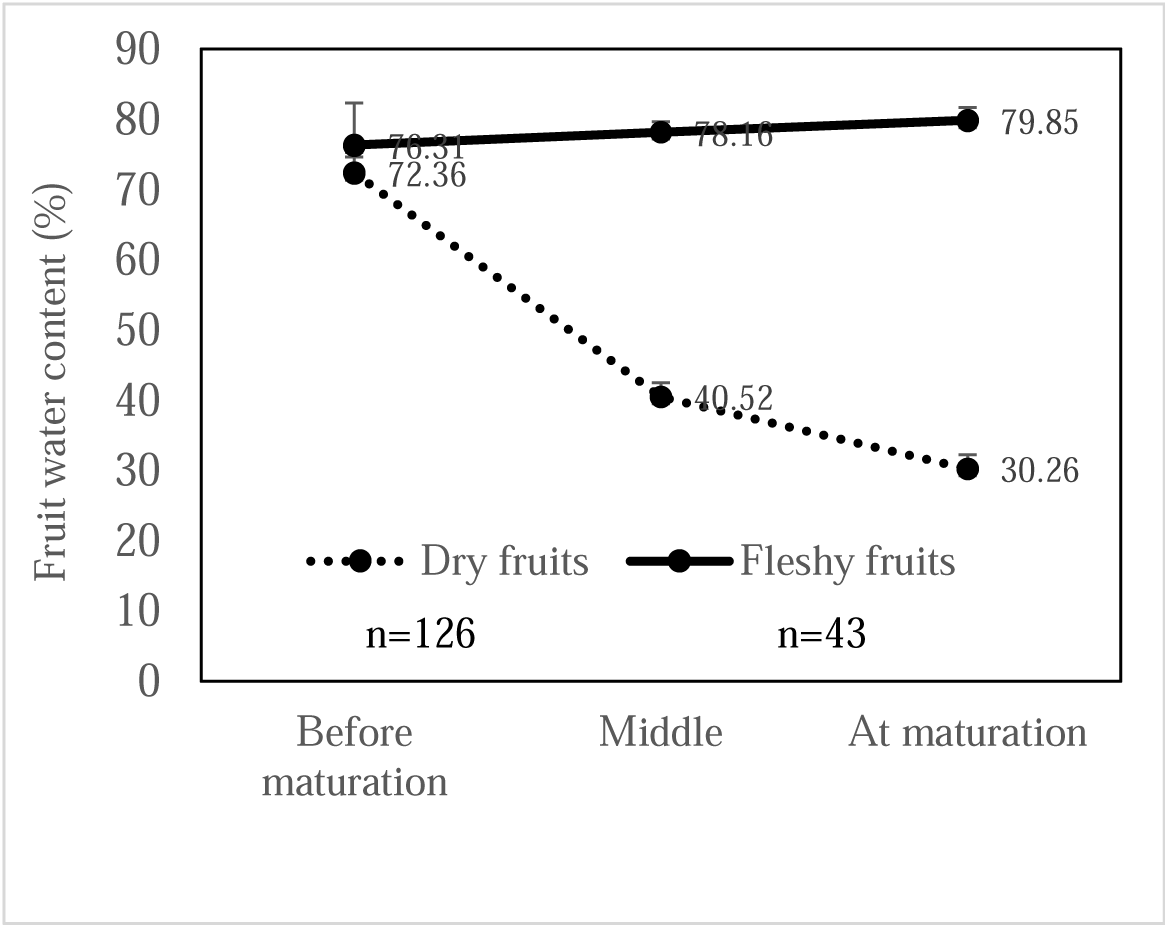
Water content shift trend chart of fleshy and dry fruits from juvenile to maturation

### 3.3 Water content shift patterns of fruits after maturity

Based on speed of water loss, two general modes were found: rapid and slow (Fig. 4). Dry fruits exhibit rapid water loss, characterized by continually declining water content from the highest value at juvenile to lowest value at maturity, followed by rapid water dissipation after maturity, or then sudden decrease of water content for some dehisced fruits. In contrast, fleshy mature fruits have slow rates of water loss (Fig. 4).

**Fig. 4.**
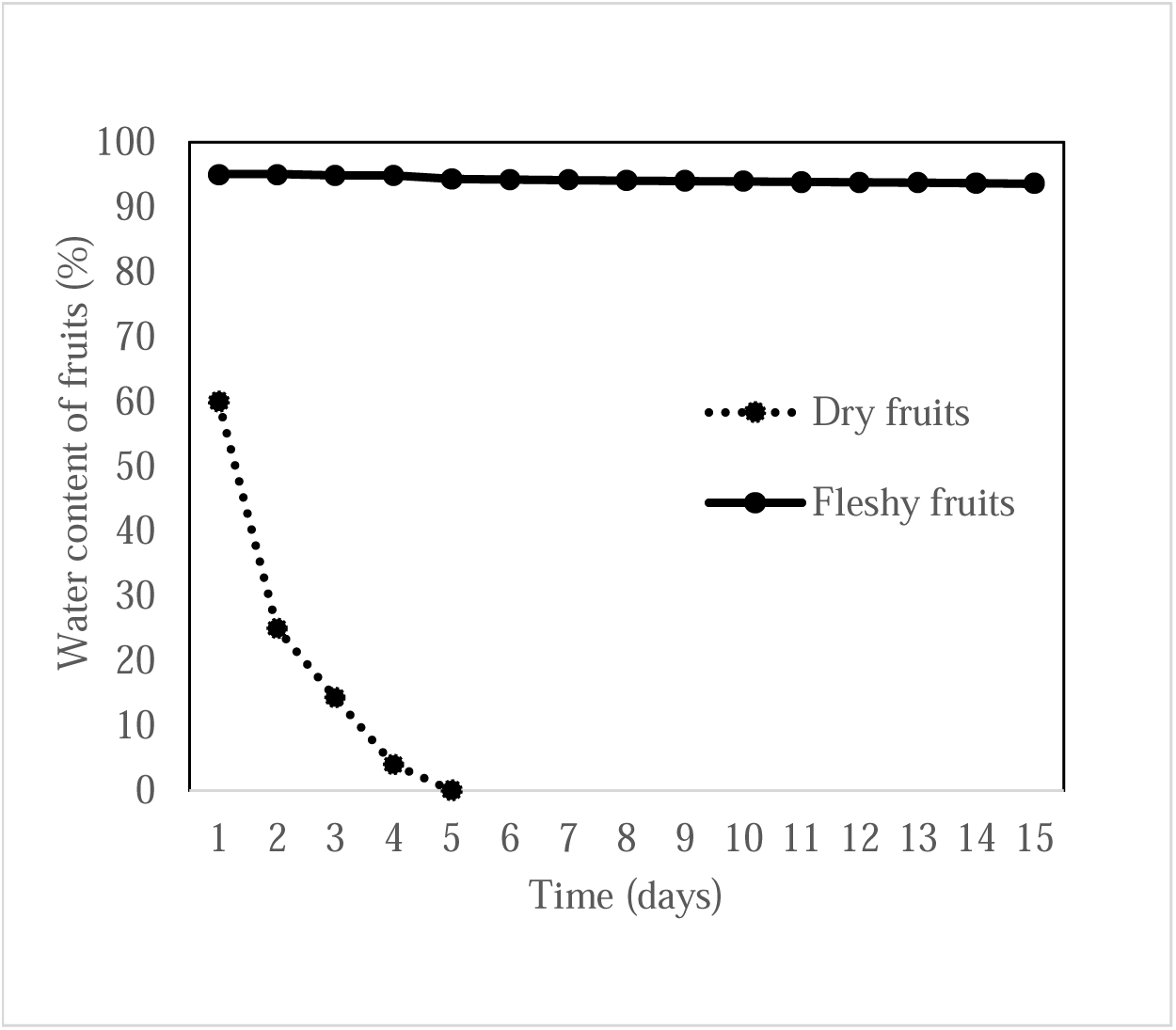
Water content shift with time for typical fleshy and dry fruits after maturation (sketch map)

### 3.4 Fruit drying period and their effect factors

There was a huge variation in fruit water content and length of the drying period among the study species (Fig. s2). For instance, fruits of *Cocos nucifera* (a type of drupes), weighing over 3kg, were not dried after 6 months of air drying.

Significant correlations were found between fruit drying period and other fruit variables (fresh mass, dry mass, proportion of dry mass, proportion of water, length, width, height, surface area and fruit size) (Tables 1-2, Fig. 5). Significant phylogenetic signals were present in all the variables (Table 3). The independent variables separately explained 7.9 - 48% of the variation in fruit drying time, while phylogeny explained 6 - 23% of the variation, with the exception of fruit height.

**Fig. 5.**
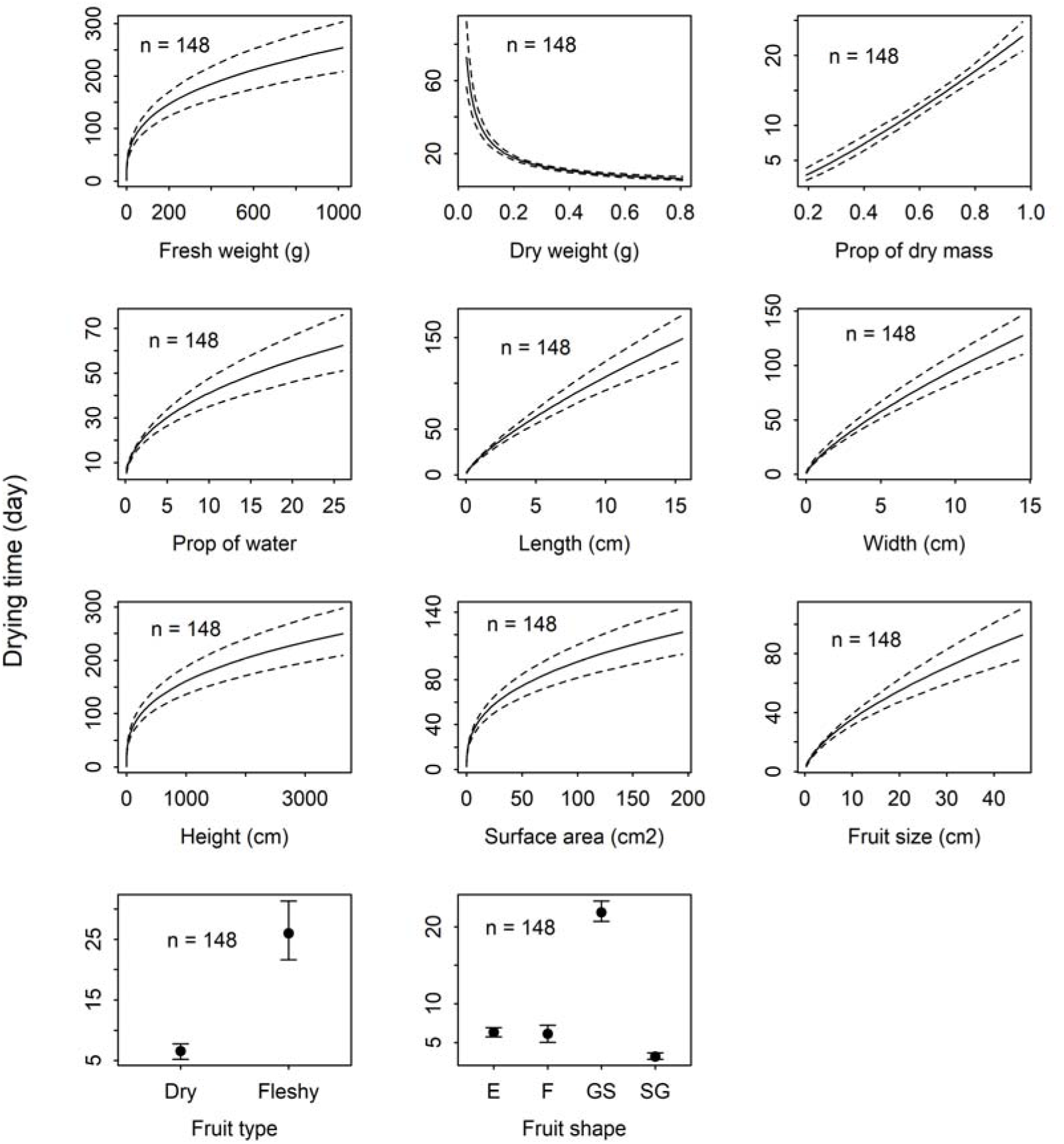
The response of fruit drying time to the independent variables such as fresh weight, dry weight, proportion of dry mass, proportion of water, length, width, height, surface area, size, type and shape of fruits.

**Table 1.**
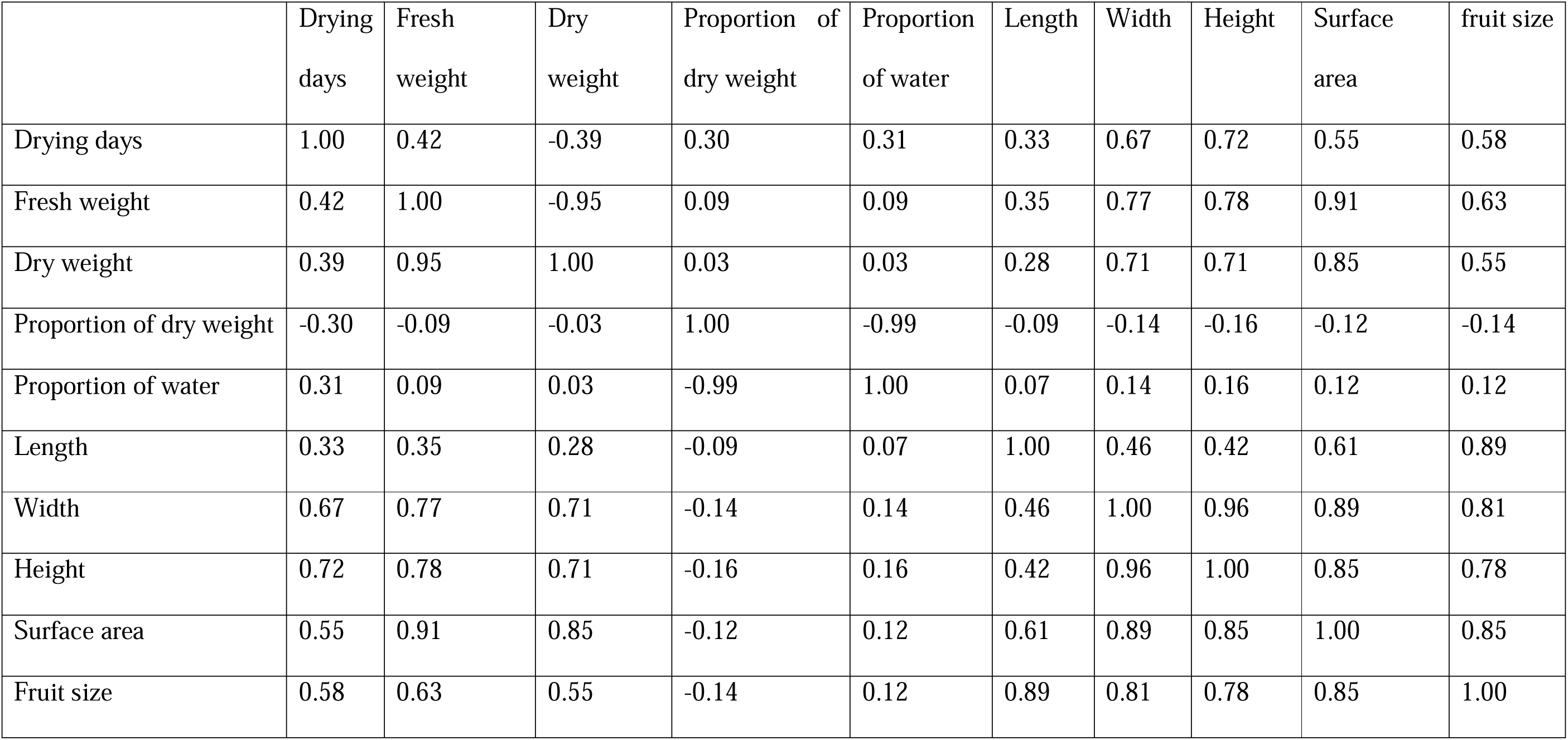
Pearson correlation coefficients of fruit drying days with variables (fruit fresh weight, dry weight, proportion of dry weight, proportion er, length, width, height, surface area and fruit size) in the original scale

**Table 2.** Pearson correlation coefficients of fruit drying days with variables (fruit fresh weight, dry weight, prop. of dry mass, prop. of water, height, surface area and fruit size) in the log-transformed scale

|  | Log drying<br>days | Log fresh<br>weight | Log dry<br>weight | Log proportion of<br>dry weight | Log proportion<br>of water | Log length | Log<br>width | Log<br>height | Log surface<br>area | Log fruit<br>size |
| --- | --- | --- | --- | --- | --- | --- | --- | --- | --- | --- |
| Log drying days | 1.00 | 0.73 | 0.69 | -0.36 | 0.21 | 0.39 | 0.59 | 0.72 | 0.54 | 0.58 |
| Log fresh weight | 0.73 | 1.00 | 0.97 | -0.36 | 0.17 | 0.67 | 0.75 | 0.80 | 0.79 | 0.78 |
| Log dry weight | 0.69 | 0.96 | 1.00 | -0.13 | -0.04 | 0.66 | 0.74 | 0.78 | 0.78 | 0.77 |
| Log proportion of dry weight | -0.36 | -0.36 | -0.13 | 1.00 | -0.84 | -0.19 | -0.21 | -0.25 | -0.22 | -0.22 |
| Log proportion of water | 0.21 | 0.17 | -0.04 | -0.84 | 1.00 | 0.08 | 0.06 | 0.07 | 0.08 | 0.08 |
| Log length | 0.39 | 0.67 | 0.66 | -0.19 | 0.08 | 1.00 | 0.59 | 0.49 | 0.91 | 0.93 |
| Log width | 0.59 | 0.75 | 0.74 | -0.21 | 0.06 | 0.59 | 1.00 | 0.82 | 0.88 | 0.79 |
| Log height | 0.72 | 0.80 | 0.78 | -0.25 | 0.07 | -0.49 | 0.82 | 1.00 | 0.72 | 0.69 |
| Log surface area | 0.54 | 0.79 | 0.78 | -0.22 | 0.08 | 0.91 | 0.88 | 0.72 | 1.00 | 0.96 |
| Log fruit size | 0.58 | 0.78 | 0.77 | -0.22 | 0.08 | 0.93 | 0.78 | 0.69 | 0.96 | 1.00 |

**Table 3.** Mutual information (MI) between fruit types or fruit shapes and other variables.

|  |  | Log drying<br>days | Log fresh<br>weight | Log dry<br>weight | Log<br>proportion of<br>dry weight | Log<br>proportion of<br>water | Log<br>length | Log<br>width | Log<br>height | Log surface<br>area | Log fruit<br>size |
| --- | --- | --- | --- | --- | --- | --- | --- | --- | --- | --- | --- |
| Fruit<br>types | MI | 2.492178e-01 | 0.074 | 0.03 | 0.091 | 0.069 | 0.035 | 0.059 | 1.20777<br>5e-01 | 0.045 | 0.034 |
|  | *p*-value | 2.495366e-07 | 0.089 | 0.126 | 0.005 | 0.234 | 0.260 | 0.025 | 1.45836<br>6e-04 | 0.154 | 0.252 |
| Fruit<br>shapes | MI | 3.528987e-01 | 0.186 | 0.183 | 0.097 | 0.098 | 0.143 | 0.145 | 2.43206<br>7e-01 | 0.140 | 0.112 |
|  | *p*-value | 2.177122e-08 | 0.006 | 0.003 | 0.051 | 0.054 | 0.012 | 0.002 | 3.35375<br>6e-06 | 0.013 | 0.035 |

### 3.5 Correlations between relative and absolute speed of fruit water loss and fruit types and shapes

Fruit type, shape and surface area were found to have positively significant relationships with rate of fresh fruit decrease in mass on the first drying day (Fig. 6, Fig. s3), and their contributions were 27.39%, 20.60% and 16.00%, respectively. Significant phylogenetic signals were observed in all the variables. The independent variables separately explained 21% to 49% of the variation in rate of mass loss of fruits, while phylogeny explained 3.8% to 4.4% of the variation.

**Fig. 6.**
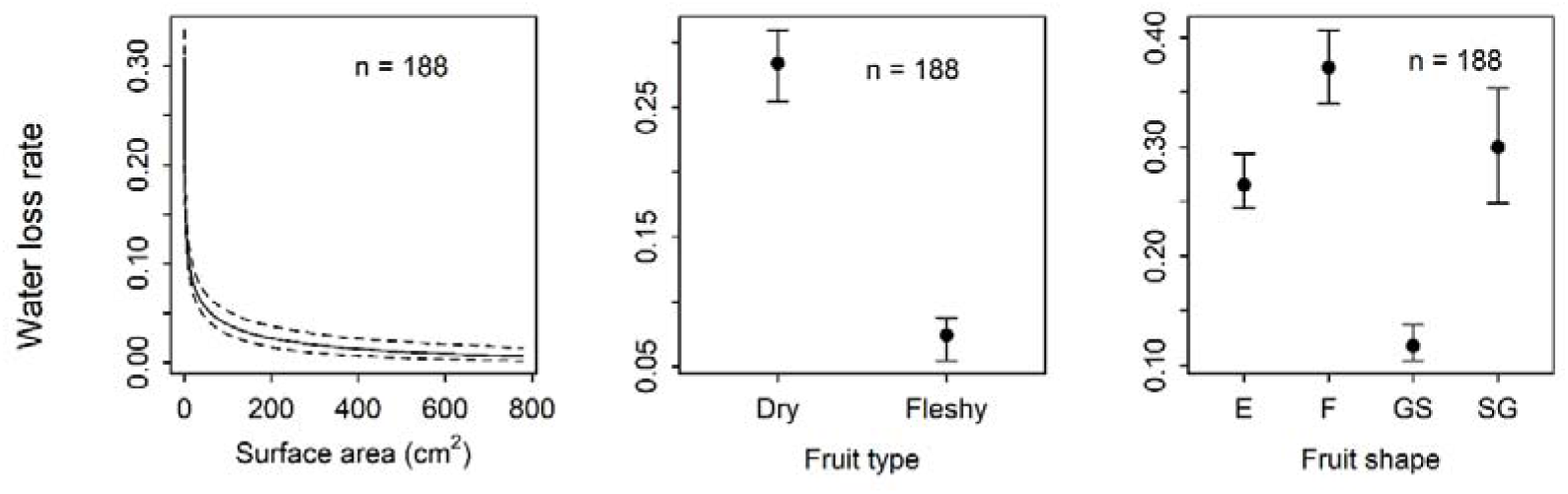
Response of relative water loss rate to the independent variables such as fruit surface, types and fruit shapes.

Dry fruits in the late stages of development stage had a significantly higher rate of absolute water loss than fleshy fruits (Fig. 7). The rates of absolute water loss vary across different fruit developmental stages. The species with smooth and bright surfaces of fleshy fruits at fruit maturity (e.g., species in Caprifoliaceae, Solanaceae, Rosaceae, Cornaceae) have the lowest water permeability of pericarp. For dry fruits, water permeability of the pericarp increases from the young to mature fruit stage, whereas for fleshy fruits water permeability of pericarp remains basically unchanged.

**Fig. 7.**
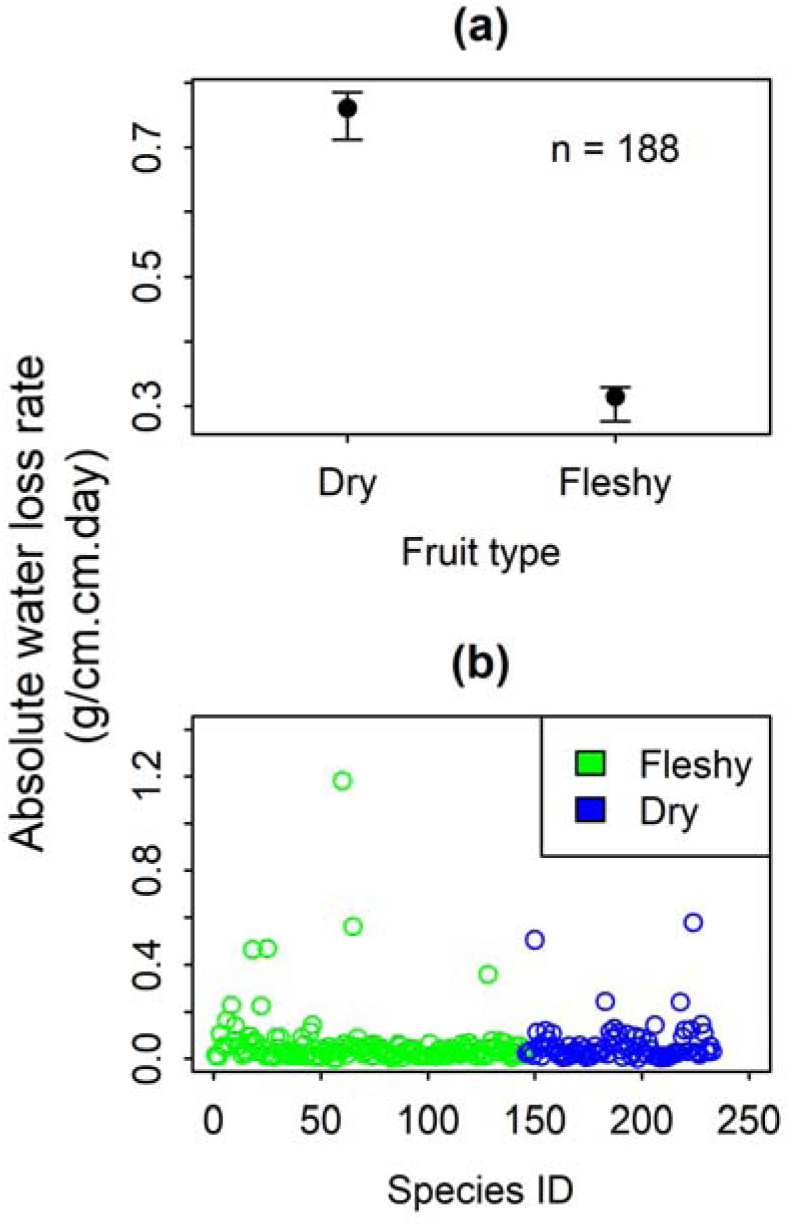
Absolute water loss rate for dry and fleshy fruits

## Discussion

### Water loss rate of fruits, fruit shapes and other several responsible factors

Our fruit drying experiment showed that mature fleshy fruits had a longer period of high-water content than dry fruits after water supply of fruits was cut off. Several biotic factors may be responsible for this. First, the shapes of fleshy fruits tend to be spherical or nearly globular (Fig. 2d), and globular spheres have the smallest surface area per unit weight. Second, we found that a higher fruit water content correlates with longer fruit freshness or fleshiness (Fig. 5). In this study, the water content of fleshy fruits reached its maximum when fruits were ripe, whereas that of dry fruits was very low at ripeness (Fig. 2). Third, the duration of fruit drying period is positively related to fruit height, and fleshy fruits generally have greater weight than dry fruits (Fig. s4) (5, 9, 17), delaying fleshiness of fleshy or dry fruits. Lastly, fleshy fruits often possess lower water permeability of pericarp than dry propagules in later development or ripening stages (Fig. 7) due to less lignin and more waxes or thicker cuticles in the pericarp of the fleshy fruits (Fig. 1) (31).

Being originated from open habitats (8), species with dry fruits have developed a series of strategies to facilitate propagule dispersal by wind, including appropriate forms (such as elongated or flaky shapes, and other forms with wing-like extensions or flanges to increase surface area for water evaporation or propagule dispersion), small fruit mass, low water content at fruit maturity, fragile pericarp (in dehiscent fruits), and high permeability pericarp near or at fruit maturity (Fig. 7). As a result of long-term adaptation strategy, fleshy fruits attract birds and mammals due to their succulent pericarp, which has high water content and abundant nutrition (large propagule mass) (5, 9, 17). In addition, origin of fleshy or dry fruit shapes controlled ultimately by genes (22) may be related to shaded versus open habitats, respectively, due to role of wind (for instance, wind velocity, wind direction and their time of duration may affect origin and evolution of fruit shapes, named as the fruit shape origin hypothesis, another topic that deserve further attention in the future) or drought (21).

### Fruit shapes and water content shifts during fruit development

The water content of dry fruits attached to the maternal plant decreased progressively from the juvenile stage to maturity under continuous water supply to fruits (Fig. 2). In contrast, water content of the majority of fleshy fruit types (for instances, berries, pomes, pepos and hesperidium etc.) increased slightly up to the point of ripeness, peaking at maturity or keeping higher water content throughout development. These findings suggest distinct patterns of water content variation during fruit development for dry and fleshy fruits.

Fruit shape is considered as one of the key factors (the other is pericarp permeability discussed as following) behind the above patterns of water content shift, due to allometric expansion between fruit surface area, podetium thickness and propagule mass. Based on Corner’s rule, stem/branch sizes should be positively related to their attached structures, such as leaves, flowers and propagules (23–25). As an apparatus of mechanical support and transport water, podetium (a kind of twig) thickness (or cross-sectional areas of fruit pedicels) was positively related to propagule mass (17). During fruit development, both cross section of the podetium and fruit mass would enlarge, however, surface areas of elongated or flaky dry fruits increase faster than that of spherical or nearly globular fleshy fruits in the case of equal weight.

If we assume that fleshiness of fleshy fruits results from water balance, then dryness of dry fruits can be attributed to water imbalance in water supplement and loss. As dry fruits develop, the amount of water supplied by fruit pedicels is less than that lost through the surface area of fruits, leading to a gradual decrease in water content for dry fruits due to the accelerated expansion of the surface area of the non-spherical fruits.

The water content values (*C_i_*, in %) of circular dry fruits can be expressed by following equation to (see methods):

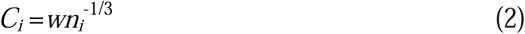

Where *w* is water content (%) of the fruits during the initial young stage and *n_i_* is multiple by which the fruit mass increases. As *n_i_* value increases with fruit growth, water content *C_i_* will decline. This is one of the fundamental causes of decreasing water content in non-spherical dry fruit types. In this study, the data in several fruit developmental stages observed from a few of species (in Ulmaceae) proved the correctness of this model.

For dry fruits with various shapes (non-spherical), the above model may be transformed to following equation:

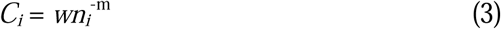

Where *m* (0<*m<*1) is shape coefficient of dry fruits with various shapes. Of course, the formula *C_i_* = *wn_i_*^-m^ holds true when the permeability of the fruit cuticles is consistent across developmental stages. However, permeability of the fruit cuticles varies across different stages of fruit development for dry fruits, and the above model may be transformed again to following equation:

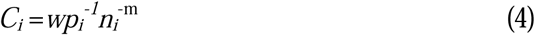

Where *p_i_* (*p_i_* >1, for dry fruits) is defined as water variation coefficient of the pericarp related to permeability, *p_i_* values will increase as dry fruits develop. For fleshy fruits, *p_i_* values are generally equal or approximate to 1. For berries, drupes, pepos, pomes and hesperidiums, *p_i_* values can be less than 1 and its minimum value should be about 0.8 based on our observational data. This variation can help explain why water content of globular dry fruits declines as they become mature.

In fact, the water inside the fleshy fruits is also unbalanced. The rising water content for majority of fleshy fruit types reflects that the input of water to the fruits is greater than the output during fruit development due to lower pericarp permeability.

However, as for plant species with drupes and rosehips, the increase or decrease pattern of the fruit water content depend on proportion of seed mass relative to the total fruit mass. If the proportion of seed mass in fruits is sufficiently large, the pattern follows a decline. For instances, in *Prunus trilobata*, where the proportion exceeds 80%, the water content declines as the seed mass (which has lower water content at maturity, 56.70 ± 3.51%) becomes dominant. Additionally, the higher water content observed in the mature fruits of *Magnolia denudata* and *Magnolia biondii* (both with follicles, a kind of dry fruits) can be attributed to their higher fruit mass, lower cuticular permeability and thicker pericarp.

### Pericarp permeability variation of fleshy and dry fruits and their environmental mechanism

Our results showed that fleshy fruit pericarps not only display a low relative rate of water loss but also an absolute low loss of water in later development stages of fruits (The values for dry fruits vary with ripeness, sample times or water content of fruits) (Fig. 7). Low water loss indicates that the pericarp of fleshy fruits exhibits low permeability due to pericarp waxification (or less lignin content), which may be attributed to long term evolution to weak light in shaded habitats (31–32) (Fig. 1). Correspondingly, strong light in the open habitats may lead to an evolution of higher pericarp permeability of dry fruits in the late stage of fruit development due to lignification occurring (being beneficial for water transportation) in pericarp (31–32) (Fig. 1), resulting in a hard shell on the outer layer of dry fruits. However, not all fleshy-fruited plants originate from shaded habitats, for example, origin process of cheliophytes with drupes and rosehips may be more complex in evolutionary history, and their lignification occurred merely in seed coats, leading to hard exotestal (Fig. 1).

Moreover, low water loss indicates that the pericarp of fleshy fruits exhibits low permeability due to thicker cuticles or more wax content in pericarps, which may evolve due to the lower temperature on the surface of the fruits in shaded habitats than in the open for a long time. Lower temperature in shaded habitats favor the accumulation of waxes in cuticles, sustaining lower cuticle permeability and plentiful water in fleshy fruits (26–27). The slow water loss rate in fleshy fruits initially suggests that structure and amounts of waxes in the cuticle of pericarps serve as a major barrier against water loss from fruits (29–29). Variations in temperature among plant organs are related to their thickness, thermal capacity and configuration (30). Under the same external conditions and radiation, fleshy fruits (global shapes) generally show the lowest temperature variation compared to dry fruits (elongated or flaky shapes) due to their thickness, which is primarily determined by radiation time and is also related to fruit size and maturity (30). Therefore, cuticles of global or spherical dry fruits that originated in open habitats have more lignin and less wax, leading to higher permeability than fleshy fruits evolved from shaded habitats.

Additionally, water acts as an adhesive to some extent, helping to keep fruit bodies intact in shaded habitats. The declining water content for mature dry fruits may bring about scarcity of water and wax (or other organic compounds) on fruit surface, increasing their pericarp permeability or causing pericarp cacking (33).

Studies on angiosperm phylogeny have shown that fleshy fruits (globular or spheroidal shapes, small globular shapes) arose in shaded habitats and disappeared in open habitats many times (8, 14, 17), suggesting that changes in lignin and wax content (or other wax traits) or cuticle thickness on the surface of fruits may play a key role due to habitat change or vegetation dynamics driven by climate (7) (see Fig. 1). Consequently, their water availability modes would transform, for instance, dry-fruited species, like *Persicaria perfoliata* and *Carex baccans*, which have lived in shaded habitats for an extended period, may evolve into species with freshy fruits (8, 14) owing to lignin content reduction and wax content increment in pericarp of fruits induced by shaded environment according to the punctuated equilibrium theory (34, 35). Nearly all mature dry fruit pericarps (such as the capsules of Liliaceae,

Orchidaceae and Cucurbitaceae) of sciophytes were found to show a tendency toward waxification and fleshiness (i.e., higher water content). Conversely, fleshy-fruited plants such as *Vitex negundo* var. *heterophylla* and *Cotinus coggygria* var. *cinereus*, which have lived in open habitats for an extended period may exhibit a transition from fleshy to dry fruit types during fruit development due to seed coat lignification and small fruit sizes.

These indicate that fruit types, displaying significant phylogenetic signals (Fig. 2a, c), are long-time evolution results that distributing parallel habitats, favoring the species to form convergent traits (17). This can be explained by key innovations (gene variation or mutation) induced by environmental pressure, for instances, the fleshy-fruited clade was found to contain more species than its dry-fruited sister clade in tropical understory plants (16).

## Conclusions

In general, fruit type diversity is determined by fruit shape diversity and pericarp permeability variation diversity resulted from allometric growth of lignin in various organs of plants related to heterogeneous habitats. The formation of dryness for dry fruits at maturity, particularly those with elongated or flaky shapes, is attributed to water imbalance between supply and loss, driven by allometric growth relationships between fruit mass (or pedicle cross section area) and surfaces area (as constrained by Corner’s rules), combining with lignification and the increased permeability of the pericarp made by it during later stages of fruit development, especially for those with globular or spheroidal shapes due to an increase in lignin and a decline in pericarp permeability. In contrast, being a result of water balance between water supply by pedicel or stem and water loss through pericarp during fruit development, the formation of fleshy fruits is attributed to globular or spherical shapes (or greater fruit thickness), and low pericarp permeability, which is considered as a result of long-term adaption to shaded habitats for sciophytes with berries, pomes, pepos etc., or a result of long-term adaption to open habitats for cheliophytes with drupes and rosehips.

During fruit development, water content values can be modeled as *C_i_* =*wp ^-1^n* ^-m^, where *n_i_* is the number of times that the fruit weight increases, *C_i_* and *p_i_*are the fruit water content and the water variation coefficient of the pericarp related to permeability at the *n_i_-th* stage during fruit development, *w* is the initial water content of the fruits in its young stage, and *m* accounts for values affected by fruit shapes in non-spherical dry fruits.

Various fruit shapes have significant phylogenetic signals, serving as one of the key traits in the emergence of fleshiness and dryness for fleshy and dry fruits at maturity, indicating long-term adaptative evolution to heterogeneous habitats. In shaded habitats where ’ancestral’ light, temperature and wind conditions are preserved, fleshy or dry fruits have continuously originated and been lost over evolutionary time, from remote antiquity to the present. This ongoing process may be driven by factors such as fruit shapes, pericarp or testa lignin and cuticular wax, which are selected for their advantages in closed or open habitats. These factors are influenced by changes in the global vegetation canopy, resulting from climate oscillations on the Earth. Our findings provide deep insight into the ecological mechanisms behind the origins of fleshy and dry fruits, their repeated derivation across various taxonomic groups, and the evolutionary motivations for seed plants with these fruit types. In addition, fleshy fruits are likely the primitive type of fruits, because globular or spherical forms can evolve into flaky and elongated forms, whereas the reversible transformation is much more difficult.

To emphasize it, it is the correlated evolution of internal plant organs and environmental adaptations [not plant-animal interactions [that determines the fleshiness of mature fleshy fruits.

## Methods

### Site description

Water content measurement of fruits were conducted from May 2022 to October 2024 mainly in Beijing Botanical Garden of the Chinese Academy of Sciences. This study included fruits of 188 plant species (belonging to 144 genera and 72 families), in which 32 species were from Guangdong Province, Inner Mongolica, Tibet, Shandong Province and Hebei Province of China.

### Classification of fruit types and shapes

Based on water condition at fruit maturity, fruits were classified into two types: fleshy and dry. The former mainly included drupes, berries, pomes, pepos, hesperidia, rose hips and hesperidiums, and the later mainly included capsules, achenes, samaras, pods, caryopses, nuts, cremocarps, follicles and utricles.

Based on the highest values of fruit length to width ratio, length to height ratio and width to height ratio, combined with fruit diameter sizes, four fruit shape types in 29760 seed plant species (from China of Flora, 1994-2004) were classified: globular or spheroidal shapes (with any values of the three ratios < 2), elongated shapes (with two values of the three ratios > 2, and the height and width are almost equal), flaky shapes (with two values of the three ratios > 2, and values of length and width are much greater than the height) and small globular shapes (with diameter < 0.5cm).

### Water evaporation experiment of fruits - fruit water content and drying days

At least three juvenile and ripe fruits were collected randomly from individuals of the plant species. The fresh weight of the fruits was measured immediately. The fruits were then allowed to dry in open air in the laboratory, and weighed daily until they reached a constant mass. Finally, the drying time (or drying days), fresh mass, dry mass, water content, relative water loss rate (the ratio of the weight lost on the first day to the total fresh weight) and absolute speed of water loss of the fruits were calculated.

Absolute water loss rates of ripe or near ripe fruits were measured using the formula:

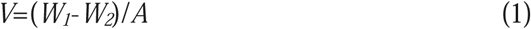

where *W_1_* and *W_2_* are the fresh weights at the start and after 1 day, respectively, and *V* and *A* are the absolute water loss rate and fruit surface area, respectively. Surface area *A* was estimated based on fruit shapes by measuring the length, width and height of the fruits. Due to the diversity and complexity of fruit shapes, the surface area was calculated and adjusted using the appropriate formula and specific coefficients.

The determination of fruit maturity is mainly based on the changes in color of the fruit skin, combined with the phenological period (15, 36). Based on this standard, the fruit development stages such as before or after fruit ripening, were determined.

### Statistical analysis

#### Fruit types and shapes

We built a phylogenetic tree for all the species (29760 in total) using the R package V.PhyloMaker2 (37). Two traits were considered in our study, including fruit types and fruit shapes. Both of them are categorical. For fruit types, we considered two categories: fleshy and dry. For fruit shapes, we considered four categories: globular or spheroidal (GS), elongated (E), flaky (F), and small globular (SG).

We conducted a measure on phylogenetic signal (δ) developed by Borges et al. (38) using their R code available at https://github.com/mrborges23/delta_statistic. The value of δ is always greater than 0. The higher the value, the stronger the phylogenetic signal between a given tree and a trait. This measure can be used for any categorical traits. For fruit types, which are binary traits, we also calculated another value (D) for phylogenetic signal, developed by Fritz & Purvis (39), using the R package caper (40). A trait with random association will have D ≈ 1, while a trait following the Brownian model will have D ≈ 0. Values of D smaller than 0 indicate that traits are more phylogenetically conserved than expected under the Brownian model, and values of D greater than 1 indicate traits that are phylogenetically over-dispersed.

We randomly selected 1000 species from the total 29760 species and extracted a sub-tree to calculate the two values, δ and D. For these sub-trees we randomly shuffled the trait values and calculated a new value for δ. For D, we randomly selected 2000 species and also performed the same procedure. All calculations were repeated 30 times.

To investigate the relationships between fruit types and fruit shapes, we built phylogenetic generalized linear models using the R package phylolm (41), which accounts for phylogenetic correlation between species. In the model, fruit types (a binary variable with a value of 1 for fleshy fruits and 0 for dry fruits) was used as the dependent variable, examining the relationships between fruit types and fruit shapes.

#### Fruits and their drying days

We also built a phylogenetic tree for the studied species using the R package V.PhyloMaker2 (36). Before conducting the following analysis, we log-transformed all the numeric variables. We tested the phylogenetic signal in drying time, fresh weight, dry weight, loss rate of fresh weight on the first day, proportion of dry mass, proportion of water, fruit length, fruit width, fruit height, fruit surface area, fruit size, fruit type and fruit shape using Pagel’s λ (42) and Blomberg’s K (43) calculated using the ‘phylosig’ function in the package ‘phytools’ v1.9-16 (43). A λ or K of 0 indicates no phylogenetic signal (44). For fruit types, we calculated the D values, and for fruit shapes, we calculated the δ vales, as described above.

We performed correlation analysis between all the variables. Since significant correlations existed between almost all pairs of variables, for numeric variables, we used the Pearson correlation coefficient, implemented in the R package stats. We conducted association analysis between a binary (fruit types) or categorical variable (fruit shapes) and a numeric variable using mutual information, implemented in the R package mpmi (45).

To investigate the relationships between drying time and other variables, we built phylogenetic linear models (PLMs) using the R package phylolm (41), which incorporates phylogenetic correlations between species. Since significant correlations existed between almost all pairs of variables, we constructed 11 models with drying time as the dependent variable and one of the 11 variables, including fresh weight, dry weight, proportion of dry mass, proportion of water, fruit length, fruit width, fruit height, fruit surface area, fruit size (the sum of length, width, and height), fruit type and fruit shape as independent variable.

To determine the contribution of each of the independent variables to the variation in drying time, we used partial R^2^ values for the PLMs (46), implemented in the R package rr2 (47). The partial R_lik_^2^ for each variable was calculated by comparing the full model (which is a PLM including the focal independent variable) with the reduced model in which the independent variable was removed, and measuring the consequent reduction in likelihood. All these models were built in the same way as described above. We also investigated the contribution of phylogeny to the variation in drying days in a similar manner. However, the reduced model was a general linear model including the corresponding independent variable without accounting for phylogeny. Additionally, PLMs were used to explore the contribution of fruit shape, type and surface area to the loss rate of fruit fresh weight on the first day in the fruit drying experiment.

#### Allometric relationships between propagule mass and surface area of circular, spherical and other forms of fruits

Taking circular and spherical fruits as examples, when propagule mass increased by a factor of *n_i_* (*n_i_*>1), surface areas of circular dry fruits would increase by *n_i_* times, while the surface areas of globular fleshy fruits would just increase by *n* ^2/3^ times (<*n*). The details of the derivation are below.

For a circular fruit, when it grows up from initial weight (*W*_1_) to a certain weight (*W*_2_), its surface area (*S*_1_, *S*_2_), volumes (*V*_1_, *V*_2_) and weights are expressed as following models:

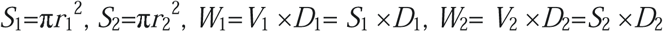

Where *r*_1_ and *r* _2_ are radii of the fruits in the two stages, *D*_1_ and *D*_2_ are densities of the fruits. If we assume *D*_1_ equals to *D*_2_, thickness is ignored (*S*_1_=*V*_1_; *S*_2_=*V*_2_), when *W*_2_=*n_i_ W*_1_, then *S*_2_= *n_i_* ×*S*_1_.

For a spherical fruit, when it grows up from initial weight (*W*_1_) to a certain weight (*W*_2_), its surface area (*S*_1_, *S*_2_), volumes (*V*_1_, *V*_2_) and weights are expressed as following models:

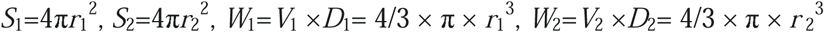

Where *r*_1_, *r* _2_, *D*_1_ and *D*_2_ are the same as the above. If we assume *D*_1_ equals to *D*_2_, when *W*_2_=*n_i_ W*_1_, then *S*_2_= *n* ^2/3^ *S*_1_.

Therefore, we suggested that the increase in surface area of fruits of other shapes is greater than that of spherical fruits as fruits develop.

## Author contributions

S.L. Yu led data collection and analysis, conceived the idea and led manuscript writing. C.R. Liu carried out phylogenetic and model analysis. Dr. H. Liu, Carol C. Baskin and X.L. Gao gave a critical revision suggestion on the draft. All authors contributed critically to the drafts and gave final approval for publication.

## Acknowledgements

This research has been supported by the National Natural Science Foundation of China (grant nos. 40771070 and 41171041) and the Beijing Natural Science Foundation (grant no. 5092015).

---

**Fig. s1.**
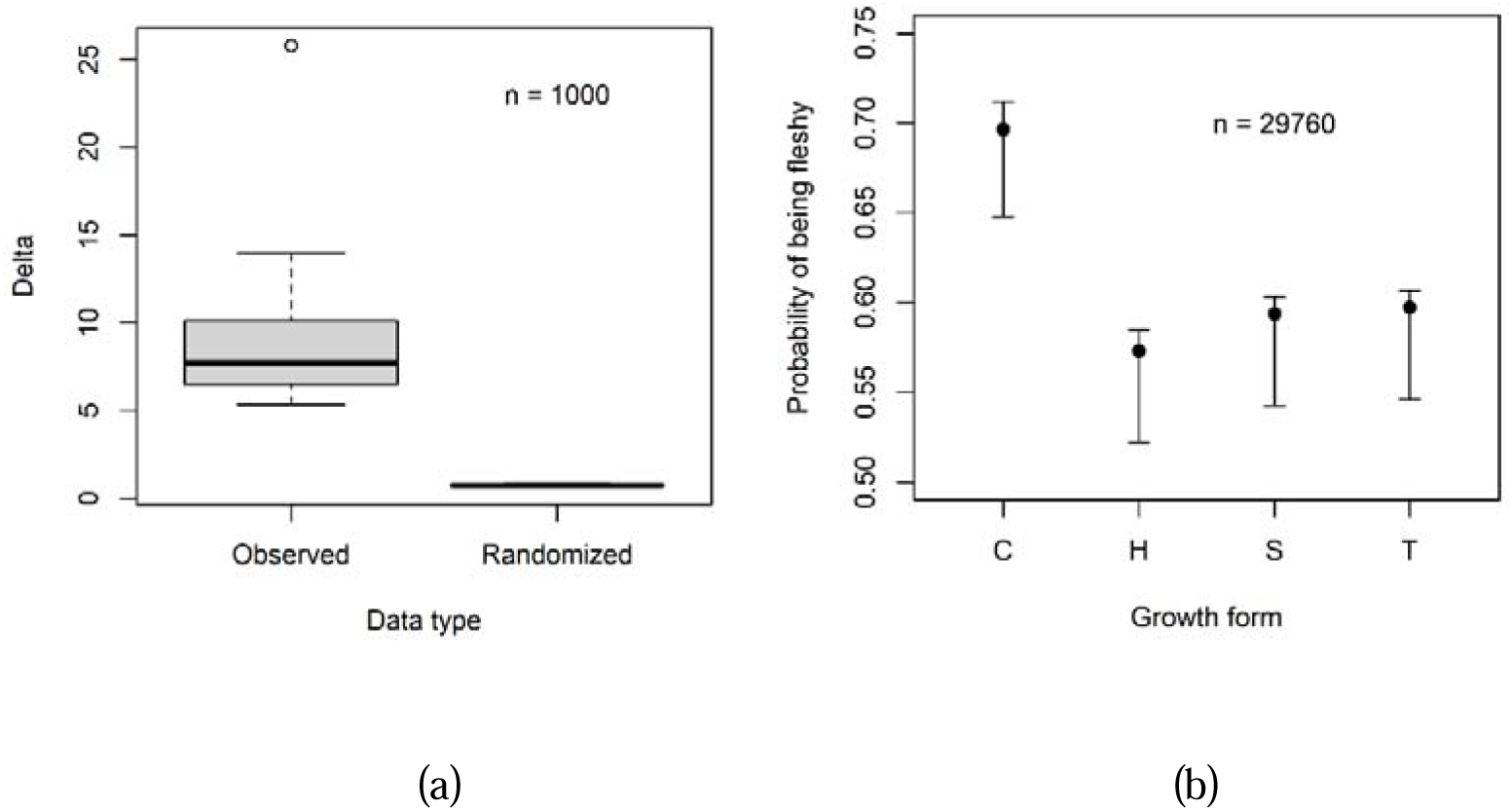
(a) Delta index measuring phylogenetic signal in growth form. Thirty sets of 1000 species were randomly selected from all the 29760 species. For each set, a value of delta (denoted as Observed here) was calculated, and the trait values were randomized and another value of delta (denoted as Random here) was calculated. (b) The probability (mean value and 95% confidence interval) that a species has a fleshy fruit for different growth form. Four categories are tree (T), shrub (S), herb (H), and climber (C).

**Fig. s2.**
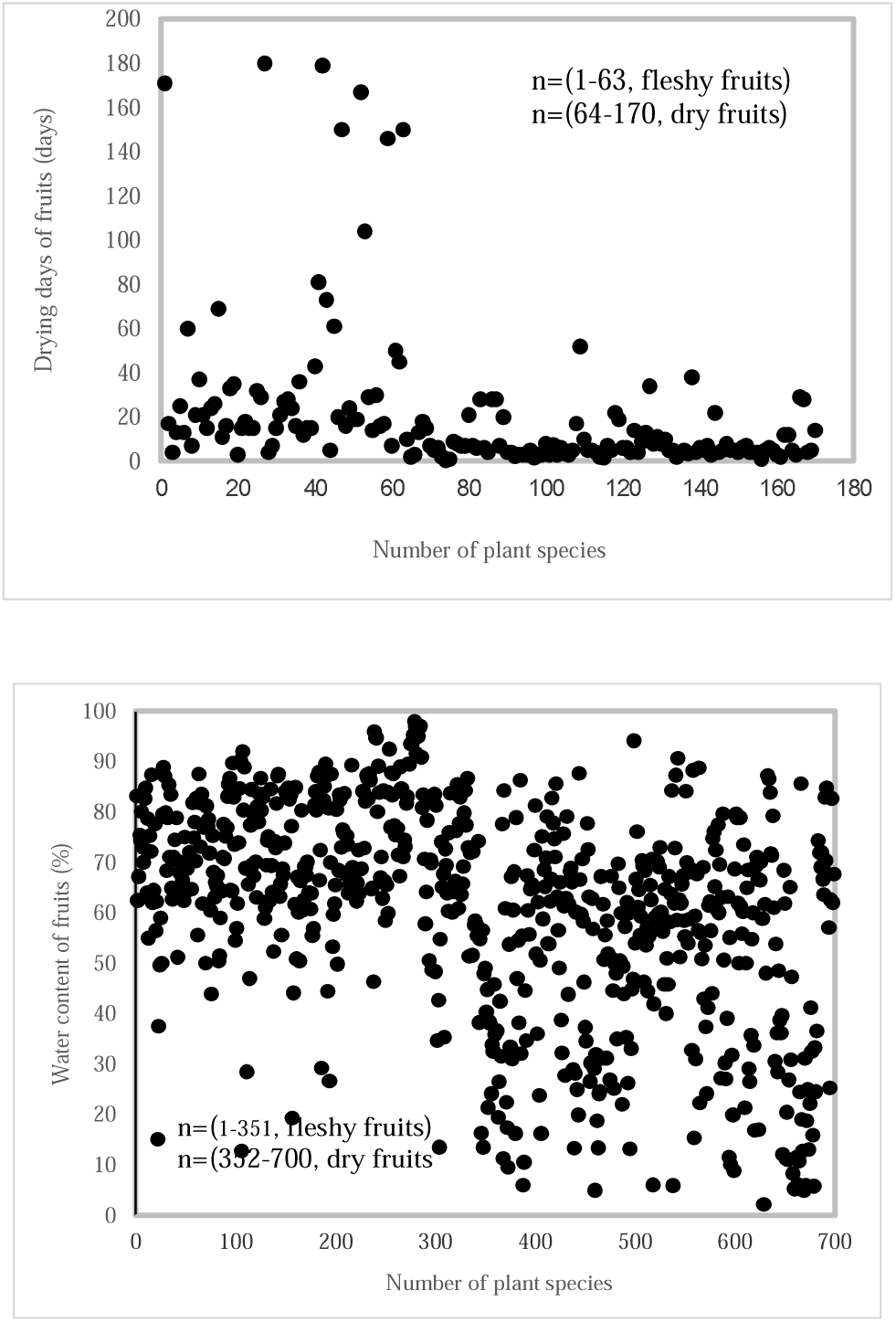
Natural drying days and water content of fleshy and dry fruits

**Fig. s3.**
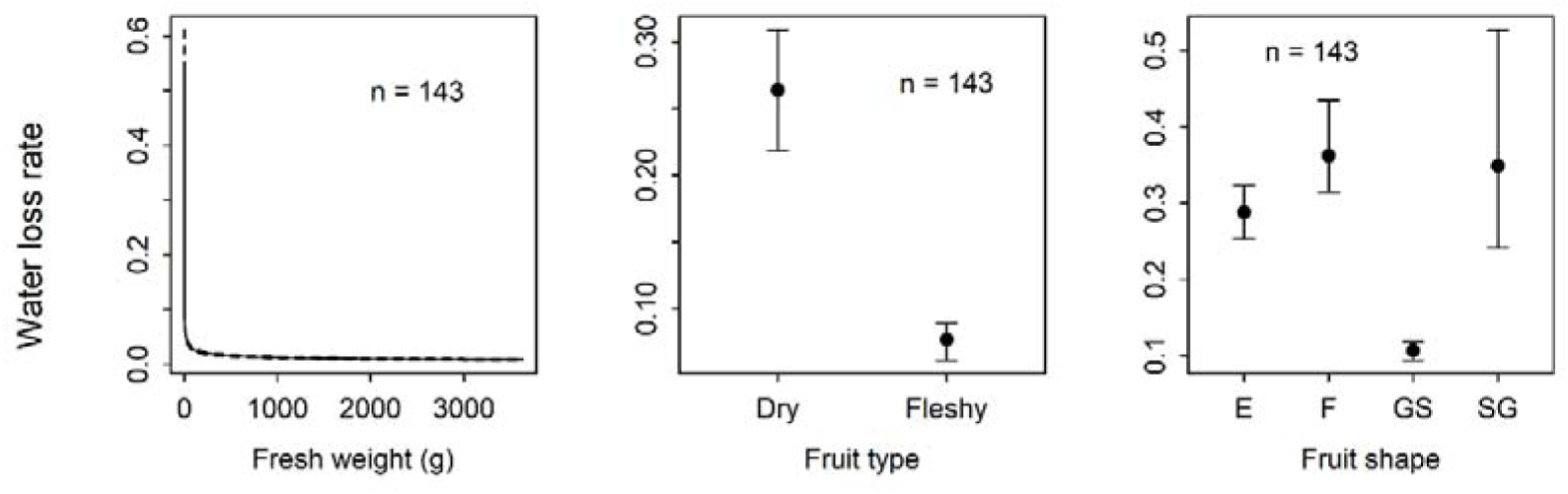
Response of water loss rate to the independent variables such as initial fresh weight, fruit types and fruit shapes.

**Fig. s4.**
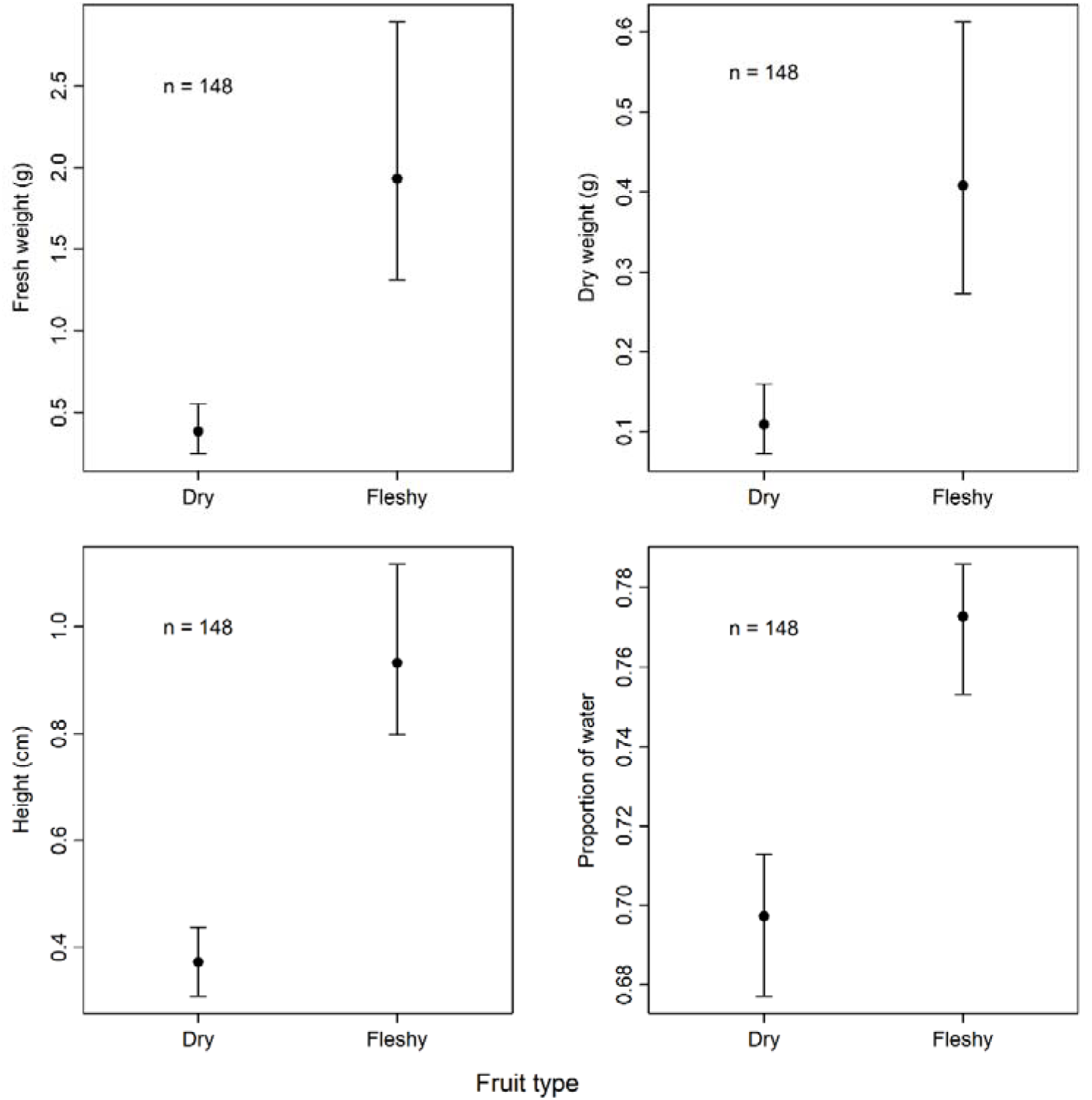
Fresh and dry weight, height (thickness), water proportion at maturation of fleshy and dry fruits.

